# Raffinose induces autophagy to promote plant growth

**DOI:** 10.1101/2024.09.02.610801

**Authors:** Sahar Magen, Sahar Daniel, Shahar Weiss, David J. Factor, Sergey Mursalimov, Yoram Soroka, Simon Michaeli, Tamar Avin-Wittenberg

## Abstract

Plant growth is governed by the integration of environmental cues and nutritional status. Under stress conditions, growth is usually attenuated in favor of stress response, creating a trade-off between growth and stress. Autophagy is a vital process in eukaryotes, maintaining cellular balance by degrading and recycling cellular components. It is triggered by various nutrient-deprivation conditions and both biotic and abiotic stresses in plants. Surprisingly, over-expressing autophagy-related genes across multiple plant species resulted in increased plant size, yield, and stress resistance, posing autophagy as a regulator of the stress-growth balance. Yet, the molecular mechanisms governing its induction remain partially understood.

In the current work, we identified raffinose-a plant-derived sugar known for its role in stress responses-as a novel plant autophagy inducer. Raffinose treatment resulted in increased biomass and yield in an autophagy-dependent manner in several plant species. We also show that raffinose activates autophagy through the SnRK1 kinase complex, independent of TOR signaling, and that raffinose treatment results in increased expression of *ATG5* and *ATG7*. We also point to possible downstream candidates operating autophagy-related biomass accumulation. Our findings offer new perspectives on the role of autophagy in maintaining a balance between plant growth and stress responses, underscoring the significance of raffinose in its regulation.

**SIGNIFICANCE STATEMENT:** The intricate balance between plant growth and stress responses is crucial for agricultural productivity, particularly as climate change intensifies environmental stressors such as drought and extreme temperatures. Usually, there is a trade-off between growth and stress response. Autophagy—a cellular recycling process essential for maintaining cellular homeostasis—plays a pivotal role in this balance. Yet, the molecular mechanisms modulating it are partially understood. Raffinose treatment enhances biomass and yields in various plant species by inducing autophagy. By elucidating the molecular mechanisms of raffinose-mediated autophagy induction, our findings provide valuable insights into potential strategies for enhancing plant resilience against climate-induced stress.

## INTRODUCTION

To survive and thrive, plants must constantly equilibrate the inherent trade-off between growth and responses to external stresses. Understanding how plants manage this delicate balance is essential not only as it is a fundamental aspect of plant life but also due to its potential effects on crop yields and food security at a time of global environmental changes. Climate change poses a severe threat to global agriculture, primarily through the increased frequency of extreme weather events and altered growing conditions. Rising temperatures, unpredictable rainfall patterns, and increased droughts or floods result in significant crop loss(1, 2).

One of the cellular mechanisms functioning plant yield determination, as well as plant abiotic stress response, is Macroautophagy (hereafter termed “Autophagy”)(3). It is a conserved eukaryotic degradation mechanism, breaking down cellular components, such as proteins, protein aggregates, and whole organelles in the lytic vacuole(4). Degradation targets are sequestered in double-membrane autophagosomes, which travel to the vacuole, and their content is recycled for reuse(4, 5). AuTophaGy-related (ATG) proteins function in autophagy initiation and autophagosome formation (4).

Autophagy has proven essential to many aspects of plant life. Autophagy-deficient plants (*atg* mutants) display early senescence and reduced yield(6–8). Other phenotypes include hypersensitivity to nutrient deficiency and various biotic and abiotic stresses(3, 9). Interestingly, overexpression of *ATG* genes in the model plant *Arabidopsis thaliana* (Arabidopsis) resulted in increased yield and abiotic stress resistance(10). Similar findings were observed in other plant species(11–13), suggesting that autophagy may modulate the interplay between stress and growth.

Two kinase complexes play a crucial role in integrating environmental signals to balance growth and stress responses, both of which impact autophagy: SnRK1, which activates autophagy(14, 15), and TORC, which inhibits it(16). Overexpression of the SnRK1 catalytic subunit KIN10 in Arabidopsis led to constitutive activation of autophagy, and a *kin10* knockout mutant failed to activate autophagy(15, 17). Conversely, overexpression of TOR blocks activation of autophagy by nutrient starvation, salt, and osmotic stresses, while autophagy induced by oxidative and ER stress is unaffected. Thus, autophagy activation can be TOR-dependent or independent(16). It, therefore, seems that various environmental conditions activate autophagy through distinct mechanisms, integrating the outputs of both TOR and SnRK signaling pathways.

Sugars are a primary cellular carbon source but can also mediate growth in a signaling/regulatory capacity. One example is trehalose-6-phosphate (T6P), which indicates the cell’s sucrose status and affects growth and development by regulating SnRK1 activity(18). Regulatory sugars have been shown to induce autophagy in metazoans(19), and emerging evidence indicates a similar function in plants(20). A study of the resurrection plant *Tripogon loliiformis* demonstrated that trehalose accumulates during its desiccation, inducing autophagy(21). This finding was strengthened by a recent publication, in which inhibition of trehalose degradation in maize resulted in autophagy activation and increased plant biomass(22). Moreover, sugar accumulation has been observed in autophagy-impaired plants(23, 24). These data indicate that other sugars might be involved in plant autophagy induction.

Raffinose is part of the raffinose family oligosaccharides (RFOs), which are α-1,6-galactosyl extensions of sucrose. The galactosyl moiety is donated by galactinol(25). It is used as a transportable sugar in several plant species and stored in seeds(26, 27). Raffinose functions in plant stress-response, accumulating in leaves during abiotic stress, specifically drought(28, 29), and also in the base of leaves during developmental senescence(30). Moreover, overexpression of raffinose synthesis enhances plant drought tolerance(31). Several mechanisms have been proposed regarding the function of raffinose in abiotic stress response: maintaining cell membrane integrity, functioning as an osmoprotectant, and stabilizing photosystem II(32). These mechanisms highlight raffinose as a structural component of resistance mechanisms, yet its possible role in signaling has not been suggested. In metazoans, raffinose was shown to induce autophagy in HaCaT cells(19, 33). In addition, raffinose accumulated in leaves of Arabidopsis *atg* mutants under nitrogen replete and deplete conditions(34) and in green fruit of tomato *ATG4*-KD plants(23).

In this study, we uncover raffinose as a novel inducer of autophagy in plants, operating through an SnRK1-mediated, TOR-independent pathway. Our findings show that raffinose treatment not only stimulates autophagy but also enhances plant growth and yield in an autophagy-dependent manner. We highlight a new role for raffinose in plants and expand our understanding of autophagy regulation at the intersection of plant growth and stress response.

## RESULTS

### Raffinose treatment induces autophagy in Arabidopsis plants

Raffinose was previously shown to induce autophagy in mammalian cells(19). We thus wished to examine this function in a plant system. First, we performed a GFP-release assay to assess metabolic flux. ATG8 is a ubiquitin-like protein anchored to the autophagosome membrane(5). The fusion protein GFP-ATG8 is used as an autophagosome marker. In addition, since the free GFP moiety is more stable than ATG8 in the acidic conditions of the vacuole, the release of free GFP from the fusion protein can be used as a measure of autophagic flux(35). We sprayed the foliage of GFP-ATG8a Arabidopsis plants with solutions containing raffinose at various concentrations supplemented by tween20 as a surfactant. We collected whole rosettes and examined the ratio between GFP-ATG8a and free GFP by Western blot. Increased GFP release following raffinose treatment was observed, suggesting amplified autophagic flux in these plants (Figure 1A,B). The amount of GFP released was dependent on the raffinose concentration used, with 0.8mM yielding the highest ratio between free GFP and GFP-ATG8a. We also examined the formation of GFP puncta by confocal microscopy in GFP-ATG8a leaves treated with or without raffinose. As in the GFP-release assay, a significantly higher number of GFP puncta, indicative of autophagosomes, was observed in raffinose-treated plants compared to the control (Figure 1C,D).

**Figure 1:**
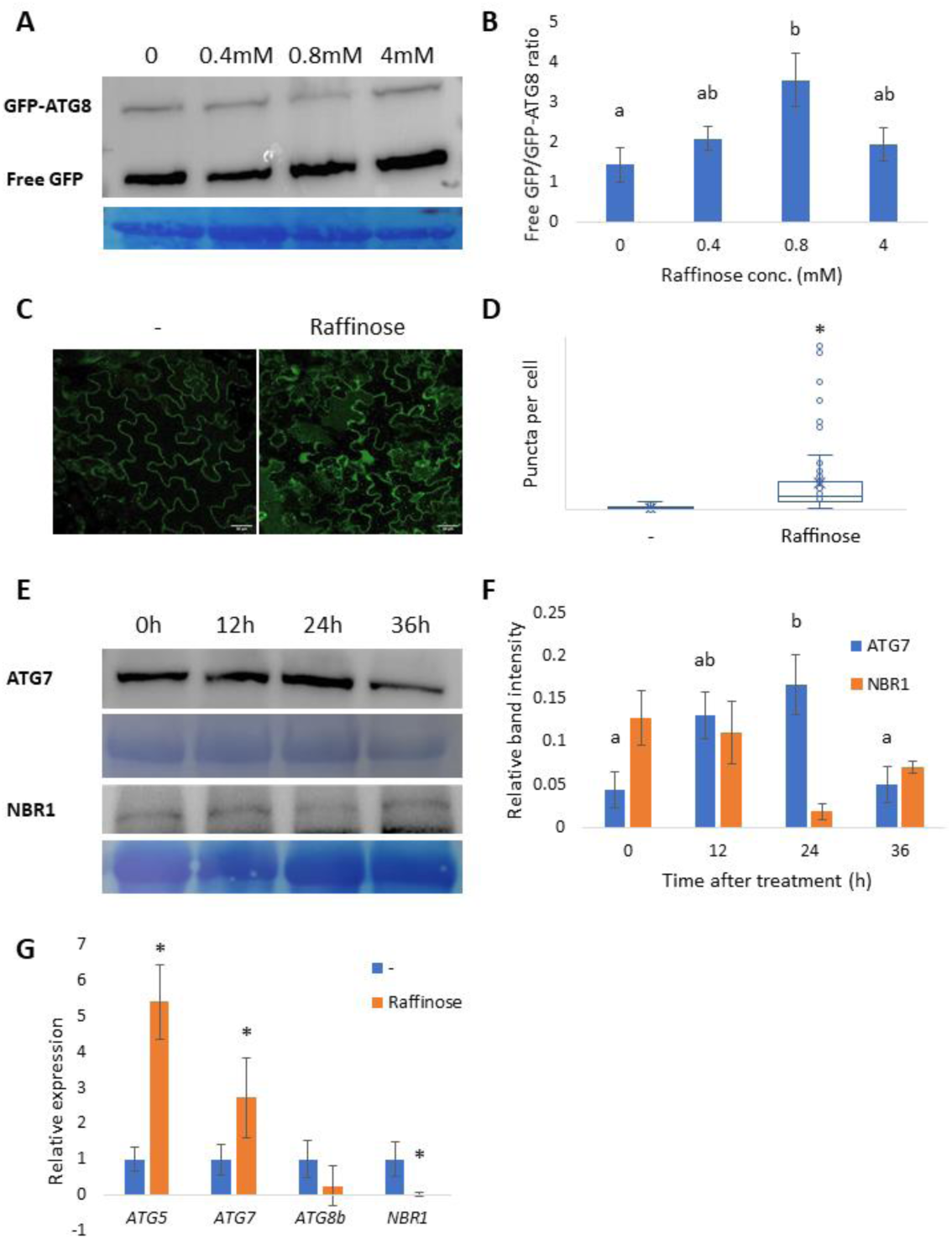
Raffinose induces autophagy in Arabidopsis leaves. A,B: Arabidopsis plants expressing GFP-ATG8a were germinated on 1% sucrose Nitcsh plates for two weeks and transferred to pots in short-day conditions. After two more weeks, the plants were sprayed once a week with a solution containing raffinose at different concentrations (0,0.4, 0.8, 4mM) and 0.05% tween20 as a surfactant. The rosette was harvested after four sprays, total proteins were extracted and examined by Western blot using anti-GFP antibody. (A) Representative image of the gel. Top panel, anti-GFP antibody, bottom panel, Coomassie blue staining. (B) Free GFP/GFP-ATG8 ratio as quantified by ImageJ. Data are presented as mean ± SE. Letters denote statistical differences following ANOVA and Tukey-Kramer test (p<0.05, n=3). C,D: Arabidopsis plants expressing GFP-ATG8a were germinated on 1% sucrose Nitsch plates for two weeks and transferred to pots in short-day conditions. The plants were grown for four additional weeks and sprayed with a solution containing two raffinose concentrations (−0mM, Raffinose – 0.8mM) with 0.05% tween20. Leaf samples were collected 18 hours after spraying and treated with 100μM Concanamycin A for autophagosome visualization. Samples were imaged by confocal microscopy (C), and the number of GFP puncta was quantified (D). Data are presented as a box & whiskers plot. An asterisk denotes statistical differences between treatments in the WT following Tukey-Kramer test (p<0.01,n=80). E;F: WT Arabidopsis plants were germinated on 1% sucrose Nitsch plates for two weeks and transferred to pots in short-day conditions. The plants were grown for 4 additional weeks and sprayed with 0.8mM Raffinose solution with 0.05% tween20. Leaf samples were collected at 0,12,24 and 36h after spraying; total proteins were extracted and examined by Western blot using an anti –ATG7 and anti-NBR1 antibodies. (E) Representative images of the gels. Bottom panel for each protein, Coomassie blue staining. (F) Protein band intensity normalized by protein as quantified by ImageJ. Data are presented as mean ± SE. Letters denote statistical differences following ANOVA and Tukey-Kramer test (p<0.05, ATG7 – n=4, NBR1 - n=3). No significant difference was observed for the NBR1 samples. (G) WT Arabidopsis plants were germinated on 1% sucrose Nitsch plates for two weeks and transferred to pots in short-day conditions. The plants were grown for 4 additional weeks and sprayed with solutions containing no sugars (-) or 1mM Raffinose with 0.05% tween20. Leaf samples were collected 24h after spraying; total RNA was extracted, and gene expression of selected *ATG* genes and *NBR1* was examined by qRT-PCR. Data are presented as mean ± SE; asterisk denotes significant differences between treatments in student’s *t-test* (p<0.05, n=4-6).

To pinpoint the timing of autophagy activation by raffinose, we performed a time-scale experiment. Wild-type (WT) plants were sprayed with raffinose, and their foliage was collected at various time points following treatment. We examined the protein amounts of ATG7 and NBR1, functioning in autophagosome formation and selective target degradation, respectively(5). ATG7 protein levels increased following raffinose treatment, peaking at 24h post-treatment, and decreased to initial protein levels after 36h (Figure 1E, top panel, Figure 1F). Interstingly, NBR1 levels displayed the opposite trend to ATG7, with the lowest levels at 24h post-treatment (Figure 1E, bottom panel, Figure 1F). Increased levels of ATG7 and reduced levels of NBR1, possibly suggesting amplified degradation, are indicative of higher autophagy activity(10), corresponding with our GFP-ATG8a results.

We also examined the expression of *ATG* genes and *NBR1* after raffinose treatment. We sprayed WT plants with solutions containing no sugars or raffinose, collected leaf samples after 24h, and analyzed gene expression by q-RT PCR (Figure 1G). The RNA levels of *ATG5* and *ATG7* significantly increase, corresponding to increased ATG7 protein levels (Figure 1E,F). Surprisingly, *NBR1* RNA levels were reduced (Figure 1G), again corresponding with the observed NBR1 protein levels (Figure 1E,F). Thus, we cannot determine whether the reduction in NBR1 protein amounts results from degradation, reduced expression, or both.

### Raffinose treatment increases plant growth and yield in an autophagy-dependent manner

Trangenic over-expression of *ATG* genes in Arabidopsis and rice increased plant biomass and yield (10, 12). We thus checked whether raffinose treatment resulted in similar outcomes. WT Arabidopsis plants were sprayed with solutions containing raffinose at different concentrations, supplemented with tween20 as a surfactant. Raffinose treatment resulted in larger rosettes (Figure 2A), and greater fresh and dry weight (Figure 2B, Sup. Figure 1A). The effect was dependent on the concentration of raffinose used, and hormesis was displayed, with higher raffinose concentrations resulting in a milder size increase. We determined 0.8mM or 1mM as the working concentration used in further experiments. Interestingly, adding tween20 to the solution resulted in lower raffinose concentrations required for biomass increase (Sup. Figure 1B), indicating that adding a surfactant to the solutions aids raffinose penetration to the leaf tissue.

**Figure 2:**
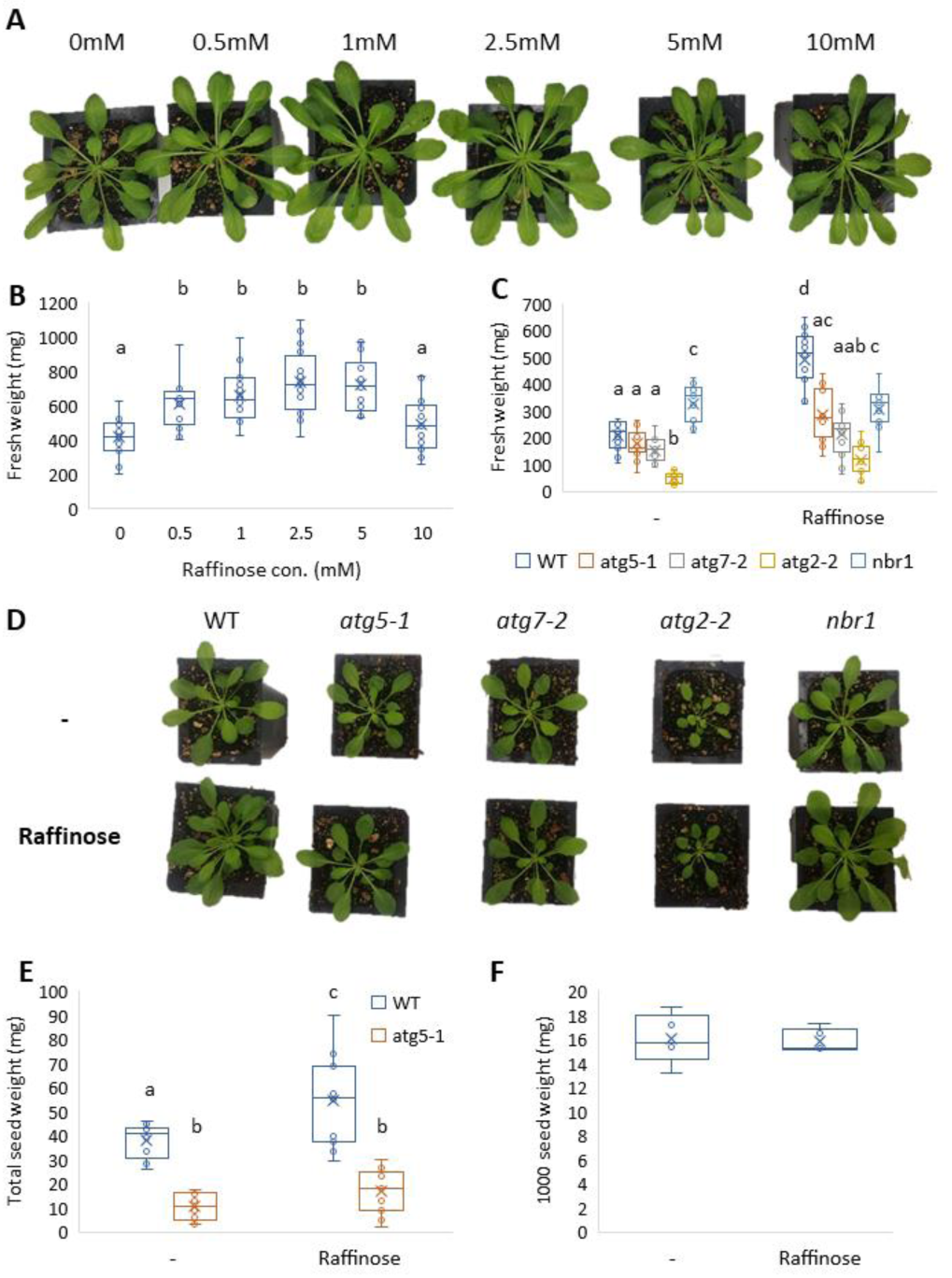
Raffinose application positively affects plant biomass and yield in an autophagy-dependent manner. A,B. WT Arabidopsis plants were germinated on 1% sucrose Nitsch plates for two weeks and transferred to pots in short-day conditions. After one more week, their foliage was sprayed every week for four weeks with raffinose solution in various concentrations (0,0.5, 1, 2.5, 5,10mM) with 0.05% tween20 as a surfactant. One week after the last spray, entire rosettes were collected. (A) Representative image of the plants. (B) Rosette fresh weight. Data are represented as box&whiskers plot. Letters denote statistical differences between treatments in the WT following ANOVA and Tukey-Kramer test (p<0.05, n=13-16). C,D. WT, *atg5-1, atg7-2, atg2-2 and nbr1* mutant Arabidopsis plants were germinated on 1% sucrose Nitsch plates for two weeks and transferred to pots in short-day conditions. After one more week, their foliage was sprayed every week for four weeks with raffinose solution in two concentrations (-: 0, raffinose: 1mM) with 0.05% tween20. One week after the last spray, entire rosettes were collected. (C) Rosette fresh weight. Data are represented as box&whiskers plot. Letters denote statistical differences between treatments in the WT following ANOVA and Tukey-Kramer test (p<0.05, n=16). (D) Representative image of the plants. E,F. WT and *atg5-1* Arabidopsis plants were germinated on 1% sucrose Nitsch plates for two weeks and transferred to pots in long-day conditions. After one more week, their foliage was sprayed once a week for four weeks with two raffinose concentrations (-: 0, Raffinose: 1mM) with 0.05% tween20. Plants were grown for seeds, and total seeds were collected per plant (E) Total seed weight per plant. Data are presented as box&whiskers plot. Letters denote statistical differences between treatments in the WT following ANOVA and Tukey-Kramer test (p<0.05, n=12). (F) Weight of 1000 seeds from treated and non-treated WT plants. No significant difference was detected between the treatments following student’s *t-test* (n=5).

Next, we wanted to ascertain whether the observed biomass accumulation after raffinose treatment resulted from autophagy activation. WT and *atg* mutant plants (*atg5-1*, *atg7-2*, *atg2-2*, and *nbr1*) were sprayed with solutions containing no sugars or 1mM raffinose, and their rosette fresh and dry weight was measured. As expected, WT plants displayed a significantly larger biomass following raffinose treatment. However, no effect was seen for the *atg* mutants (Figure 2C, Sup. Figure 1C), suggesting that raffinose-mediated plant growth is autophagy-dependent. We then measured the total seed yield of WT and *atg5-1* mutants sprayed with solutions with and without raffinose. As previously described, untreated *atg5-1* mutants had reduced seed yield compared to WT plants (7, 10, 36). Raffinose treatment resulted in higher total seed weight in WT but not in *atg5-1* plants (Figure 2E). We also measured the Thousand Seed Weight of raffinose-treated and untreated WT plants. No significant difference was observed (Figure 2F). We thus postulate the difference in total seed yield stems from higher seed quantity rather than larger seed size. Taken together, our findings suggest that raffinose treatment induces autophagy to promote plant biomass and yield accumulation.

Raffinose is one of several RFOs(32). We thus wished to examine whether other RFOs or substances functioning in raffinose biosynthesis (Figure 3A) can also achieve the same biomass increase observed following raffinose treatment. We treated WT plants with raffinose, raffinose precursors, or other RFOs (galactinol and stachyose) and examined their fresh weight. As expected, raffinose treatment resulted in increased biomass. Interstingly, galactinol, but not stachyose treatment, resulted in a similar increase in biomass (Figure 3B). To verify that the biomass increase is autophagy-dependent, we repeated the experiment with *atg5-1* mutants. We also checked that the increase in plant biomass stems specifically from raffinose treatment and not from a general addition of exogenous carbon by treating the plants with 3mM glucose solution (Sup. Figure 2). The use of a higher glucose concentration accounted for the amount of carbon in a glucose molecule, which is a monosaccharide, compared to raffinose, a trisaccharide. As observed in Figure 3B, raffinose and galactinol promoted an increase in plant biomass, while *atg5-1* plants were unaffected. Treatment with 3mM glucose did not influence plant biomass, suggesting that the increase in biomass stems from a specific function of raffinose rather than the general addition of carbon.

**Figure 3:**
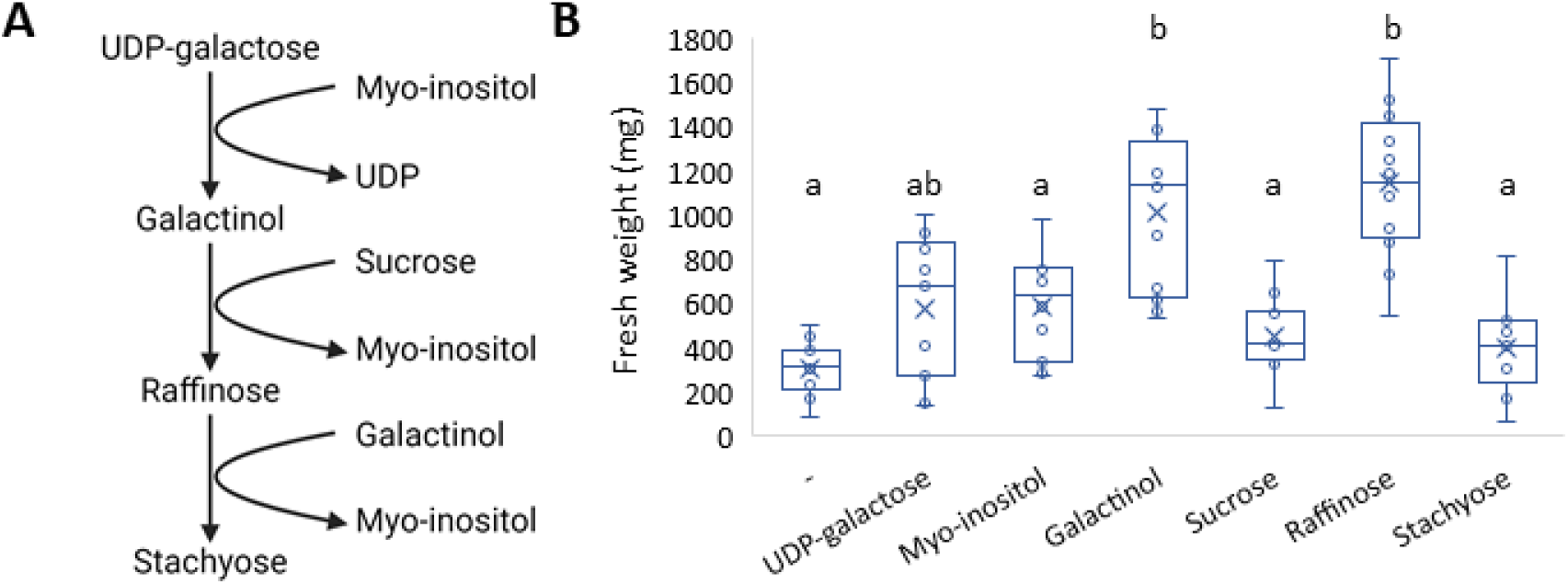
Galactinol treatment has a similar effect on plant biomass as raffinose. (A) Simplified scheme of raffinose biosynthesis adapted from Yan et al . (B) WT Arabidopsis plants were germinated on 1% sucrose Nitsch plates for two weeks and transferred to pots in short-day conditions. After one more week, their foliage was sprayed every week for four weeks with solutions containing so sugars (-) or 1mM UDP-galactsoe, myo-inositol, galactinol, sucrose, raffinose or stachyose with 0.05% tween20 as a surfactant. One week after the last spray, entire rosettes were collected and rosette fresh weight measured. Data are represented as box&whiskers plot. Letters denote statistical differences between treatments in the WT following ANOVA and Tukey-Kramer test (p<0.05, n=9-12).

### Raffinose increases plant biomass via SnRK1 in a TOR-independent manner

Autophagy function is regulated by two kinases: TORC, inhibiting autophagy(16), and SnRK1, activating it(14, 15). SnRK1 can induce autophagy directly, in a TOR-independent manner, or indirectly, by inhibiting TORC(37). To assess whether these kinases are involved in raffinose-induced autophagy, we took advantage of the increased biomass phenotype observed in plants following raffinose treatment. We sprayed WT, a*kin10-1* mutants for impaired SnRK1 signaling, and TOR-OE plants for repression of autophagy by TOR with raffinose and examined their biomass after treatment. *atg5-1* was used as a negative control for raffinose response. We postulated that if raffinose functioned *via* SnRK1, inhibiting the kinase would not result in increased growth following treatment. If raffinose activated autophagy *via* TORC inactivation, the over-expression of TOR would fail to elicit an increase in biomass. As we have previously observed, WT plants exhibited increased biomass after raffinose treatment, while *atg5-1* mutants did not (Figure 4A). Interestingly, a*kin10-1* mutants did not respond to raffinose treatment, while TOR-OE plants did (Figure 4A), suggesting that raffinose activated autophagy by SnRK1 in a TOR-independent manner.

**Figure 4:**
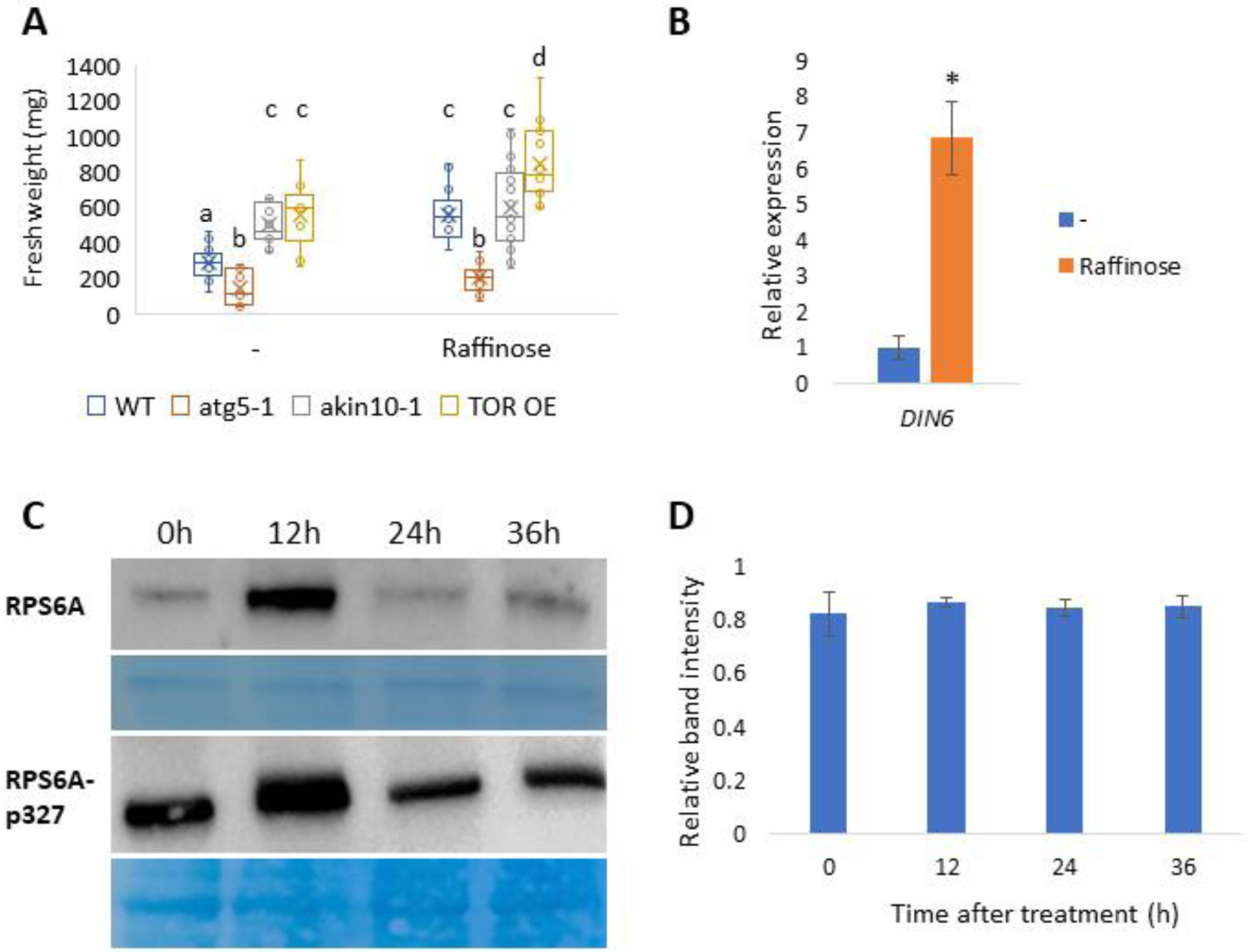
Raffinose treatment increases plant biomass in an SnRK1-dependent and TORC-independent manner. (A) WT, atg5-1, akin10-1 and TOR-OE Arabidopsis plants were germinated on 1% sucrose Nitsch plates for two weeks and transferred to pots in short-day conditions. After one more week, their foliage was sprayed every week for four weeks with solutions containing no sugars (-) or 1mM raffinose with 0.05% tween20 as a surfactant. One week after the last spray, entire rosettes were collected and rosette fresh weight measured. Data are represented as box&whiskers plot. Letters denote statistical differences between treatments in the WT following ANOVA and Tukey-Kramer test (p<0.05, n=13-16). (B) WT Arabidopsis plants were germinated on 1% sucrose Nitsch plates for two weeks and transferred to pots in short-day conditions. The plants were grown for 4 additional weeks and sprayed with solutions containing no sugars (-) or 1mM Raffinose with 0.05% tween20. Leaf samples were collected 24h after spraying; total RNA was extracted, and gene expression of *DIN6* was examined by qRT-PCR. Data are presented as mean ± SE; asterisk denotes significant differences between treatments in student’s *t-test* (p<0.05, n=6). C,D: WT Arabidopsis plants were germinated on 1% sucrose Nitsch plates for two weeks and transferred to pots in short-day conditions. The plants were grown for 4 additional weeks and sprayed with 0.8mM Raffinose solution with 0.05% tween20. Leaf samples were collected at 0,12,24 and 36h after spraying; total proteins were extracted and examined by Western blot using an anti–RPS6A and anti-RPS6Ap327 antibodies. (C) Representative images of the gels. Bottom panel for each protein, Coomassie blue staining. (D) Relative protein band intensity between RPS6Ap327 and RPS6A as quantified by ImageJ. Data are presented as mean ± SE. No significant statistical differences were found following ANOVA and Tukey-Kramer test (p<0.05, n=3).

To further confirm our observation, we examined the expression of *DIN6*, a downstream target of SnRK1(14). WT plants were sprayed with raffinose, and *DIN6* RNA levels were analyzed 24h post-treatment by qRT-PCR. DIN6 expression significantly increased in raffinose-treated plants than those sprayed without it (Figure 4B), strengthening our hypothesis that raffinose induces autophagy by SnRK1 activation. We next tested RPS6A, which is phosphorylated by S6K, a downstream target of TORC(38). We tested the protein levels of RPS6A in our time-scale experiment as well as RPS6A phosphorylated form (RPS6A-p327). We observed no difference in the ratio between RPS6A levels and its phosphorylated form (Figure 4C&D), suggesting that raffinose treatment does not alter RPS6A phosphorylation and, thus, does not affect TORC. Taken together, our results indicate that raffinose induces autophagy by activating SnRK1 in a TORC-independent manner.

### Metabolic profiling suggests a possible influence of raffinose on pectin manipulation

To identify the downstream pathways affected by raffinose treatment, we performed metabolic profiling and proteomics. For metabolic profiling, WT plants were sprayed with solutions with or without raffinose. Rosettes were collected 24h after spraying, and polar metabolites were analyzed by gas chromatography-mass spectrometry (GC-MS). We were able to annotate 25 compounds (see Supp. Table S1 for full details). Principal component analysis (PCA) revealed that non-treated samples (red circles) clustered together, while raffinose-treated samples (blue circles) were separated from them, forming a “gradient” along PC1 (Figure 5A). We attribute this variation to the treatment method. Although we sprayed the rosettes until they were completely covered in liquid, it is possible that due to differences in rosette size and leaf angles, different amounts of raffinose could penetrate the leaf, causing the differences in effect.

**Figure 5:**
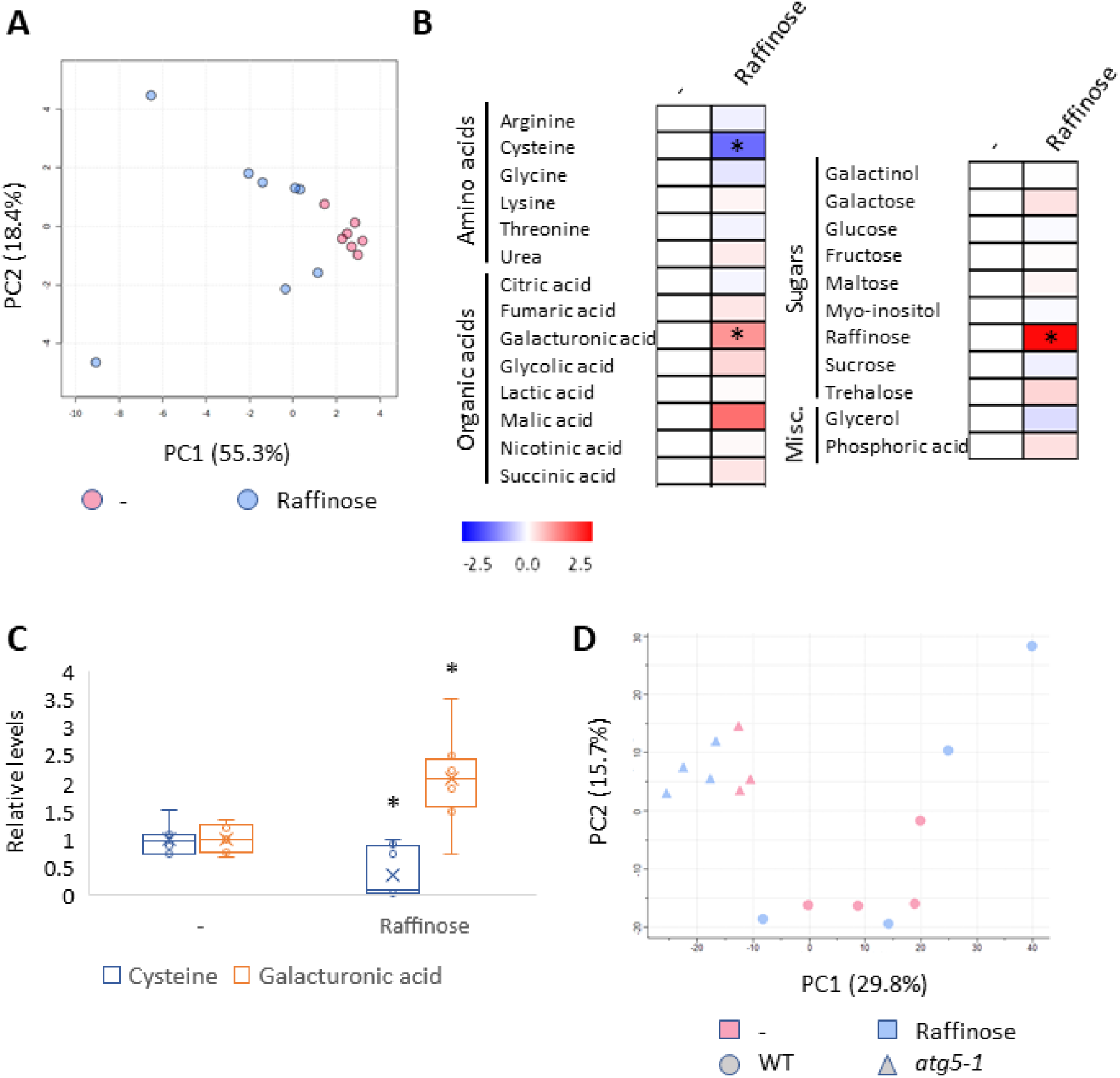
Mild changes in primary metabolism following raffinose treatment. A-C: WT Arabidopsis plants were germinated on 1% sucrose Nitsch plates for two weeks and transferred to pots in short-day conditions. The plants were grown for 4 additional weeks and sprayed with solutions containing no sugars (-) or 1mM Raffinose with 0.05% tween20. Leaf samples were collected 24h after spraying; primary metabolites were extracted and analyzed by Gas chromatography–mass spectrometry (GC-MS). (A) Principal component analysis (PCA) of primary metabolite levels. (B) Log2 values of the relative metabolic content are presented as a heat map. An asterisk denotes significant differences compared with the WT following student’s *t-test* (FDR < 0.05). (C) Relative metabolite levels of cycteine and galacturonic acid. Data are presented as box&whiskers plot; an asterisk denotes significant differences between treatments in student’s *t-test* (FDR<0.05, n=7-8). (D) WT and *atg5-1* Arabidopsis plants were germinated on 1% sucrose Nitsch plates for two weeks and transferred to pots in short-day conditions. The plants were grown for 4 additional weeks and sprayed with solutions containing no sugars (-) or 0.8mM Raffinose with 0.05% tween20. Leaf samples were collected 24h after spraying; total proteins were extracted and analyzed by Liquid chromatography–mass spectrometry (LC-MS/MS). PCA of protein levels.

We observed significant differences after raffinose treatment in three metabolites: raffinose, cysteine, and galacturonic acid (Figure 5B). Raffinose levels significantly increased, which is expected after raffinose treatment. Interestingly, levels of other raffinose precursors measured in our analysis (i.e., galactinol, sucrose, and myo-inositol) did not change after treatment (Figure 5B). Cysteine levels were reduced after raffinose treatment (Figure 5C). Autophagy activity was previously shown to be mediated by protein persulfidation(39), and it will be interesting to examine this possible connection in future research. In addition, metabolic profiling of *atg12* mutant maize seeds revealed an accumulation of cysteine in the mutants compared to WT, which the authors attributed to ROS metabolism (40). It will be interesting to examine whether raffinose treatment affects ROS metabolism *via* autophagy induction. Galacturonic acid levels increased after raffinose treatment (Figure 5C). Galacturomic acid is a major sugar residue of plant pectins(41), suggesting a possible involvement of raffinose treatment in pectin remodeling.

We also conducted a proteomic analysis of WT and *atg5-1* plants 24h after raffinose treatment (Figure 5D, see Supp. Table S2 for full details). PCA showed no separation between treated and non-treated *atg5-1* samples (blue and red triangles, respectively), yet WT and *atg5-1* samples were clearly different (circles vs. triangles). Unfortunately, we did not observe a clear separation between treated and non-treated WT samples (blue and red circles). However, we did observe a similar trend as seen in the metabolic profiling, in which the non-treated samples were closer to each other, and the treated samples were forming a “gradient” along PC1. The metabolic profiling and proteomics experiments were not performed simultaneously, suggesting this observation may stem from the treatment rather than a technical issue with a specific experiment. Interestingly, a pectin-related protein, Pectin acetylesterase 3 (PAE3, At2g46930), was slightly but significantly down-regulated in WT plants following raffinose treatment (ratio 0.7476, p-value 0.0448) but not in *atg5-1* plants. This again points to a possible connection between raffinose treatment, autophagy, and pectin remodeling.

### The effect of raffinose on plant biomass is conserved in several plant species

Enhancing autophagic activity *via* transgenic overexpression of *ATG* genes was shown to improve plant performance in the model plant Arabidopsis as well as several crop species (10, 42, 43). We thus wished to examine whether our observation that raffinose treatment increases plant biomass also translates to other plant species. We sprayed lettuce (*Lactuca sativa*), *Nicotiana benthamiana,* and wheat (*Triticum aestivum*) with solutions containing no sugars or 0.8mM raffinose and assessed the effect on plant fresh weight, dry weight, and leaf number. As observed in Arabidopsis, the plants seemed bigger (Figure 6A,C,E), and we noticed a significant increase in plant foliage fresh and dry weight in all the plant species tested (Figure 6B,D,F, Sup. Figure 3D). In addition, raffinose treatment resulted in higher leaf numbers in *Nicotiana benthamiana* and wheat (Sup. Figure 3A,B) and increased plant height in wheat (Sup. Figure 3C). As for Arabidopsis, we treated the plants with 3mM glucose as a control (Sup. Figure 2). In all plant species tested, glucose treatment did not affect plant biomass, while raffinose treatment did (Figure 6, Sup. Figure 3). Our results demonstrate that raffinose induces autophagy to promote plant growth in a specific manner.

**Figure 6:**
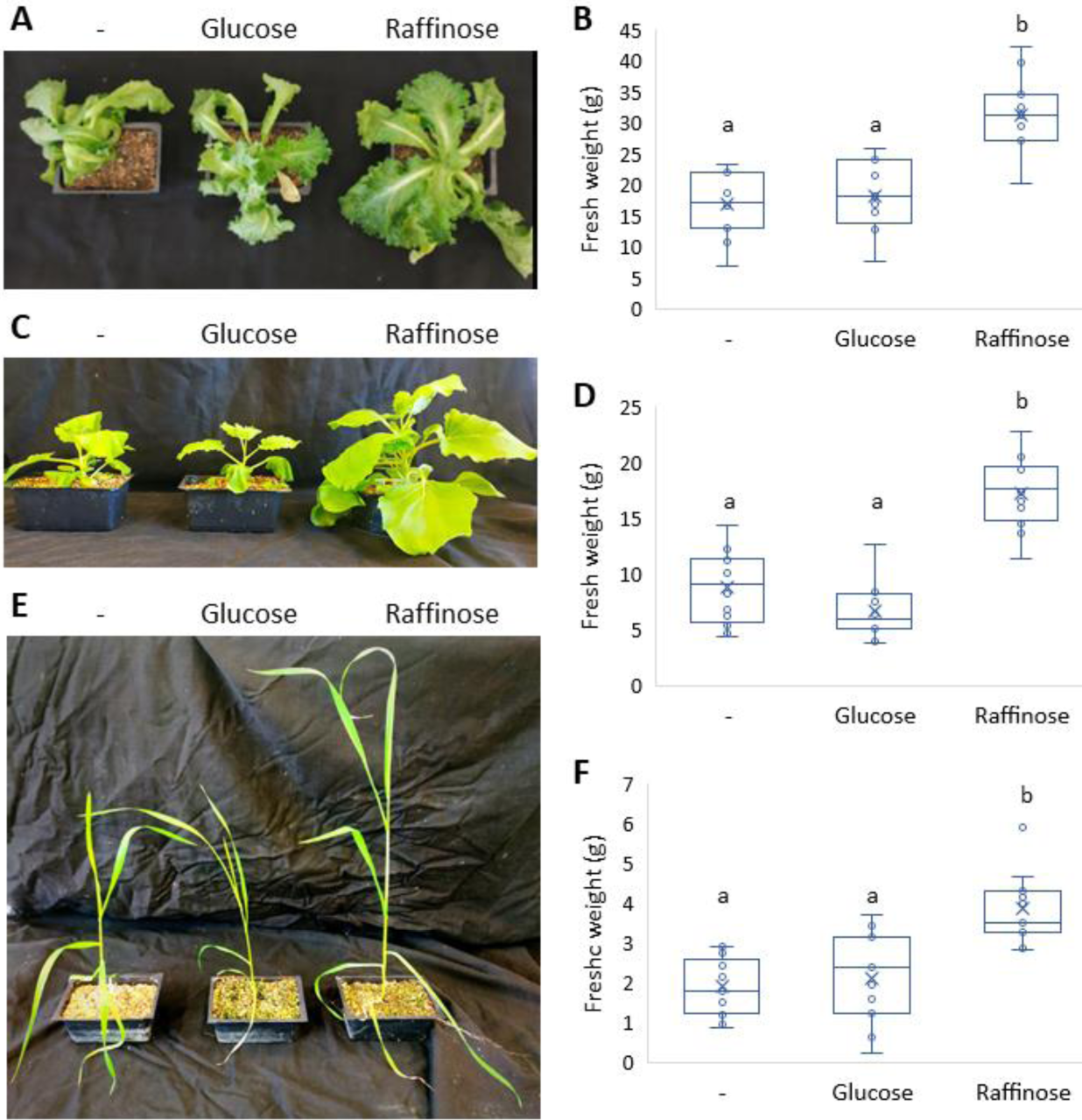
Raffinose application positively affects plant biomass in several plant species. A,B. Lettuce (*Lactuca sativa*) plants were grown in long-day conditions for 3 weeks. The plants were sprayed with solutions containing no sugars (-), 0.8mM raffinose or 3mM glucose, supplemented with 0.05% tween20 as a surfactant once a week for four weeks. A. Representative image of the plants at the end of the experiment. B. Foliage fresh weight one week after the fourth spraying. Data are presented as box&whiskers plot. Letters denote statistically significant treatments following ANOVA and post-hoc Tuckey-Kramer test (p<0.05, n=10). C,D. Tobacco (*Nicotiana benthamiana*) plants were grown in long-day conditions for 3 weeks. The plants were sprayed with solutions containing no sugars (-), 0.8mM raffinose or 3mM glucose, supplemented with 0.05% tween20 once a week for three weeks. C. Representative image of the plants at the end of the experiment. D. Foliage fresh weight one week after the third spraying. Data are presented as box&whiskers plot. Letters denote statistically significant treatments following ANOVA and post-hoc Tuckey-Kramer test (p<0.05, n=12). E,F. Wheat *(Triticum aestivum*) plants were grown in long-day conditions for 2 weeks. The plants were sprayed with solutions containing no sugars (-), 0.8mM raffinose, or 3mM glucose, supplemented with 0.05% tween20 once a week for three weeks. E. Representative image of the plants at the end of the experiment. F. Foliage fresh weight one week after the third spraying. Data are presented as box&whiskers plot. Letters denote statistically significant treatments following ANOVA and post-hoc Tuckey-Kramer test (p<0.05, n=11).

## DISCUSSION

Autophagy plays a crucial role in plant growth and development, responding to various biotic and abiotic stresses and impacting plant senescence and yield (44). Despite its significance, the mechanisms governing autophagy induction in plants remain largely unknown. In this study, we identified raffinose as a novel autophagy inducer in plants. While raffinose has been previously shown to induce autophagy in metazoans (33), it is predominantly synthesized by plants (32), suggesting that its primary roles likely evolved within the plant kingdom. Moreover, unlike the metazoan studies, which relied on cell cultures, our findings demonstrate that raffinose can trigger autophagy at the whole-plant level.

Raffinose is an abiotic stress-related sugar (32), similar to trehalose (45), another stress-associated sugar shown to induce autophagy during the desiccation of the resurrection plant *Tripogon loliiformis* (21). In maize, trehalose accumulation, achieved by chemically inhibiting its degradation, was recently found to increase plant biomass through an autophagy-dependent pathway (22). These findings align with our results, where raffinose treatment led to increased biomass in WT plants but not in *atg* mutant plants (Figure 2). We also observed that the impact on biomass accumulation is conserved in several plant species (Figure 6, Supp. Figure 3). It is intriguing that two sugars associated with stress can actually stimulate growth, which is generally suppressed under stress conditions(46). This highlights the possibility that autophagy could be an essential factor in this “decision-making” process. Similar increases in biomass and yield were also observed in Arabidopsis plants overexpressing the autophagy-related genes *ATG5* or *ATG7* (10), further strengthening our hypothesis.

It is important to note that we applied raffinose exogenously to induce autophagy and, consequently, plant biomass and yield accumulation. Thus, the physiological consequences in which raffinose activates autohpahy in plants are still unknown. As raffinose is related to abiotic stress response(32), we speculate it activates autophagy under these conditions. We also observed that the effect on plant growth was raffinose-specific and unrelated to carbon feeding, as treating plants with comparable amounts of glucose did not affect plant biomass (Figure 6, Supp. Figures 2&3). Galactinol, a raffinose precursor, also caused an increase in plant biomass following treatment (Figure 3B). Our metabolic profiling showed no difference in galactinol levels 24h after raffinose treatment (Figure 5B), suggesting that galactinol is converted to raffinose after treatment. Using exogenous application also implies we cannot rule out that other RFOs can function in raffinose-induced autophagy. It is possible that permeability constraints prevented the compounds from entering the plant and eliciting their function, thus not affecting plant growth.

How does raffinose induce autophagy in plants? Our results point to SnRK1 as the target of raffinose signaling. Mutants lacking SnRK1 activity failed to display increased biomass following raffinose treatment, and the transcript levels of *DIN6*, a downstream target of SnRK1(14), were enhanced by raffinose treatment (Figure 4A,B). Raffinose is not the only sugar influencing SnRK1 activity. Among others, T6P, a regulatory sugar promoting plant growth and development, was shown to inhibit SnRK1 activity(47). A recent publication correlated raffinose accumulation under elevated temperatures with T6P synthase activity(48). It is thus tempting to speculate that there is a possible connection between raffinose and T6P in regulating SnRK1 activity. In addition, we observed transcriptional upregulation of both *ATG5* and *ATG7* (Figure 1G). Transcriptional regulation of *ATG* genes occurs as a long-term effect of autophagy induction (4). It stands to reason that the physiological effects we observe in raffinose-treated plants (i.e. increased growth and yield) are the consequence of long-term activation of autophagy. Moreover, transgenic overexpression of *ATG5* or *ATG7* resulted in similar phenotypes (10), strengthening our hypothesis.

We still do not know how raffinose/autophagy activation causes an increase in plant biomass and yield accumulation. Our proteomics and metabolomics analyses did not result in enough significant changes to explain the observed phenotype (Figure 5A,D). We postulate this is due to sampling time, in which molecular changes have not yet been evident. Further experiments are required, collecting samples at several time points after treatment to fully understand the dynamics of the process. The metabolic profiling displayed an increase in galacturonic acid levels following treatment (Figure 5B,C), implying a change in pectin structure. Pectin is a major component of the primary cell wall, and studies have shown that modulating pectin structure can influence cell growth(49). Our observation poses pectin modulation as an interesting target for future studies regarding raffinose-induced autophagy-dependent plant growth. Further studies are needed to understand the role of autophagy in the stress-growth nexus and its regulation by raffinose.

## MATERIALS & METHODS

### Plant Material and Growth Conditions

#### Arabidopsis

*Arabidopsis thaliana* ecotype Columbia-0 was used in this study. Other lines used were p35S-GFP-ATG8a (10), *atg2-2* ((50), *atg5-1* (SAIL_129B079) (51), *atg7-2* (GK-655B06) (52), *akin10-1* (GABI_579E09) (53), and TOR-OE (Gabi_ G548(54)).

Seeds were surface sterilized with Cl2 gas for 3h, sown on Nitsch plates (Duchefa Biochemie, Haarlem, Netherlands) with 1% sucrose with 0.5% agar and imbibed for 48-72 h at 4°C in the dark. The plates were transferred for two weeks to a growth chamber (22°C at continuous light 125 μmol/m2/s and 50% rH). Seedlings were transferred to pots (25% perlite, 25% vermiculite, 50% soil) and grown under short-day conditions (8h light and 16h dark) unless indicated otherwise.

#### Additional plant species

Lettuce (Lactuca sativa), Tobacco (Nicotiana benthamiana), and Wheat (Triticum aestivum) seeds were sown in pots (25% perlite, 25% vermiculite, 50% soil) and grown in long-day conditions (16h of light and 8h of dark).

### Protein Extraction

For the time course experiment, six-week-old WT Arabidopsis plants were grown in shorth-day conditions and sprayed with solutions containing so sugras or 0.8mM raffinose (Sigma Aldrich, St. Louis, MO, USA), supplemented with 0.05% tween 20 (J.T Baker, Phillipsburg, NJ, USA). Leaf tissue was collected 0,12,24,36h after the spraying and flash-frozen in liquid nitrogen before storage at -80°C. For autophagy flux assay, six-week-old GFP-ATG8a Arabidopsis plants were grown in shorth-day conditions and sprayed with solutions containing different raffinose concentrations of (0mM, 0.4mM, 0.8mM, 4mM), all supplemented with 0.05% tween20. Leaf tissue was collected 24h after spraying and flash-frozen frozen in liquid nitrogen before storage at -80°C.

Frozen leaf tissue (∼100 mg) was ground in a bead-beater. Proteins were extracted using extraction buffer (50 mM Tris-HCl pH 7.5, 20 mM NaCl, 10% glycerol, 1% Triton, and a 1:7 ratio of protease inhibitor; 04693124001, Roche, Basel, Switzerland), based on a previously described extraction buffer (55). Samples were centrifuged for 15min at 20,000 g at 4°C, and the supernatant was collected and stored at −20 °C.

### Western blot

Total proteins were separated using a 12.5% SDS-PAGE. The proteins were transferred to a 0.2µm polyvinylidene difluoride membrane (Amersham Hybond) under a transfer buffer (56). Coomassie brilliant blue (0.25% w/v in destain solution: 50% methanol, 10% acetic acid, 40% DDW, was used to ensure equal loading. For immunodetection, rabbit anti-GFP antibody (Abcam, Cambridge, UK; AB290 1: 10,000 dilution), rabbit anti-ATG7 (Agrisera, Vännäs, Sweden; AS19 4277 1: 1,000 dilution), rabbit anti-NBR1 (Agrisera, Vännäs, Sweden; AS14 2805 1: 1,000 dilution), rabbit anti-RPS6A (Agrisera, Vännäs, Sweden; AS19 4292 1: 2,000 dilution), rabbit anti-RPS6A-P237 (Agrisera, Vännäs, Sweden; AS19 4291 1: 1,000 dilution) were used in combination with horseradish peroxidase-conjugated goat anti-rabbit antibody (GenScript, Piscataway, NJ, USA; A00098). Proteins were then imaged using the Gel Imager (Fusion FX, Vilber, Collégien, France). Protein band intensity was assessed using the ImageJ software (57).

### Confocal microscopy

Six-week-old GFP-ATG8a Arabidopsis plants grown in shorth-day conditions were sprayed with solutions containing no sugars or 0.8mM raffinose (Sigma Aldrich, St. Louis, MO, USA) supplemented with 0.05% tween 20. After 24h, individual leaves were injected with 100µM concanamycin A (ConA; Cayman Chemical Company, Ann Arbor, MI, USA), cut, and transferred to 100µM ConA solution in the dark for confocal imaging. Confocal imaging was done using the Leica SP8 CLSM system at the Volcani Institute microscopy core facility. Excitation/detection-range parameters for GFP were 488nm/500– 550nm, and emissions were collected using the system’s hybrid (Hyd) detectors. X20 dry (NA 0.75) or X63 water immersion (NA 1.3) objectives were used. Scanning was routinely done in “line” mode. The images were processed and analyzed using FIJI (ImageJ)(57).

### RNA extraction and qRT-PCR

Six-week-old WT Arabidopsis plants were grown in short-day conditions. The plants were sprayed with solutions containing no sugars or 1mM raffinose (Sigma Aldrich, St. Louis, MO, USA) supplemented with 0.05% tween 20 (J.T Baker, Phillipsburg, NJ, USA). Leaf samples were collected after 24h in 2ml tubes (6 biological replicated per treatment). The samples were flash-frozen in liquid nitrogen and stored at −80 °C until extraction.

Total RNA was isolated using TRIzol reagent (Molecular Research Center Inc. Cincinnati, OH, USA) according to the manufacturer’s instructions. Total RNA was treated with DNAse I (TURBO DNA free KIT; Invitrogen, Carlsbad, CA, USA, Dnase; Thermo Fisher Scientific, Waltham, MA, USA). The integrity of the RNA was checked on 1% (w/v) agarose gels, and the concentration was measured using a Nanodrop spectrophotometer (DeNovix, Wilmington, DE, USA). Finally, 200ng of total RNA was reverse transcribed with Superscript SentiFAST cDNA synthesis Kit (Bioline, London, UK). Real-time PCR reactions were performed in BIO-RAD HARD SHELL 96 well PCR Plates, using SYBR Green PCR Master Mix in the qRT-PCR (CFX Duet 96-well; Bio-Rad, Hercules, CA, USA). The primers used here are described in Table 1. Transcription abundance was calculated by the standard curves of each selected gene and normalized using the constitutively expressed gene ACTIN (ACT2).

**Table 1:**
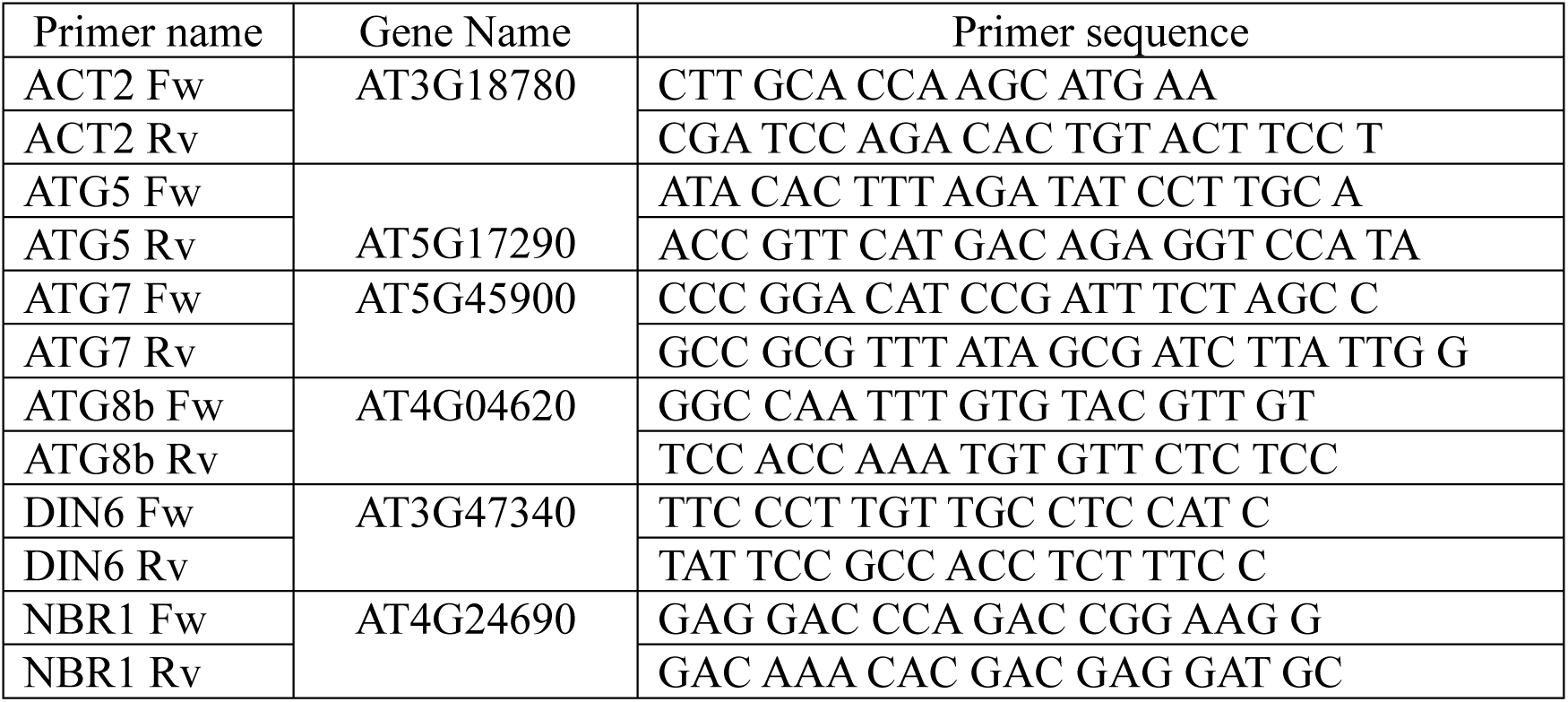
Primers used for q-PCR analysis

### Plant biomass measurement

Three-week-old WT, *atg5-1*, *atg2-2, atg7-2*, TOR-OE and *akin10-1* Arabidopsis plants were grown in short-day conditions. The plants were sprayed with solutions containing no sugars or 1mM raffinose (Sigma Aldrich, St. Louis, MO, USA) supplemented by 0.05% tween 20 (J.T Baker, Phillipsburg, NJ, USA) once a week for 4 weeks. One week after the last spray, whole rosettes were collected and weighed. For dry weight, rosettes were incubated in an oven at 60°C for 48h and weighed after drying.

### Arabidopsis yield analysis

Three-week-old WT and *atg5-1* Arabidopsis plants growth in long day conditions (16h light, 8h dark). The plants were sprayed with solutions containing no sugars or 1mM raffinose (Sigma Aldrich, St. Louis, MO, USA) supplemented by 0.05% tween 20 (J.T Baker, Phillipsburg, NJ, USA) once a week for 4 weeks. Plants were grown for seeds and floral stems were covered in paper bags to prevent seed loss. The plants were dried for one month, total seeds were collected and weighed. In addition, the weight of 1000 seeds was measured for treated and not treated WT plants.

### Treatment with raffinose precursors and RFOs

Three-week-old WT and *atg5-1* Arabidopsis plants were grown in short-day conditions. The plants were sprayed once a week for 4 weeks by solutions containing no sugars, 1mM raffinose (Sigma Aldrich, St. Louis, MO, USA), 1mM galactinol (Cayman Chemical Company, Ann Arbor, MI, USA), 1mM stachyose (Cayman Chemical Company, Ann Arbor, MI, USA), 1mM UDP-galactose (Cayman Chemical Company, Ann Arbor, MI, USA), 1mM sucrose (Duchefa Biochemie, Haarlem, Netherlands), 1mM myo-inositol (Sigma Aldrich, St. Louis, MO, USA), 1mM galactose (Sigma Aldrich, St. Louis, MO, USA) or 3mM glucose (Carl Roth, Karlsruhe, Germany), all supplemented with 0.05% tween20 (J.T Baker, Phillipsburg, NJ, USA). One week after the last spray, whole rosettes were collected and weighed.

### Raffinose treatment of additional plant species

Three-week-old Lettuce (Lactuca sativa) and Tobacco (Nicotiana benthamiana) or two-week-old Wheat (Triticum aestivum) plants were grown in long-day conditions. The plants were sprayed with solutions containing no sugars, 0.8mM raffinose (Sigma Aldrich, St. Louis, MO, USA), or 3mM glucose supplemented with 0.05% tween (20 J.T Baker, Phillipsburg, NJ, USA) once a week for 4 weeks. One week after the last spray, whole shoots were collected and weighed. In addition, the number of leaves in lettuce and the length of the blades in wheat were measured. For dry weight, shoots were incubated in an oven at 60°C for 48h and weighed after drying.

### Metabolic profiling

Metabolites were extracted following an established gas chromatography-mass spectrometry (GC-MS)-based metabolite profiling protocol (58), with slight modifications. Six-week-old WT Arabidopsis plants were grown in short-day conditions. The plants were sprayed with solutions containing no sugars or 1mM raffinose (Sigma Aldrich, St. Louis, MO, USA), supplemented with 0.05% tween 20 (J.T Baker, Phillipsburg, NJ, USA). Leaf samples were collected after 24h in 2ml tubes (7-8 biological replicated per treatment). The samples were flash-frozen in liquid nitrogen and stored at −80 °C until extraction. The samples were extracted in 700μl methanol containing an internal standard (0.2mg ribitol mL−1 water), shaken for 15min at 70°C, and centrifuged at 14,000rpm for 10min. The supernatant was transferred to new tubes, and 375μl chloroform and double-distilled water were added. Tubes were then centrifuged at 14,000rpm for 15min, and aliquots of 150μL of the upper phase of each sample were transferred to new 1.5mL tubes and dried overnight by Speedvac (Concentrator Plus, Eppendorf, Hamburg, Germany). Derivatization was carried out as previously described (58).

Polar metabolites were measured by the Agilent 7200B GC/Q-TOF. The injection and separation procedures were performed according to Dahtt using the DB-35MS column (59). Metabolite detection and annotation were performed by Quant software (Agilent, Santa Clara, CA, USA) according to an accurate mass library of known markers generated by our group and run in the same system. Following blank subtraction, the peak area of each metabolite was normalized to the internal standard (i.e., ribitol) in each sample and by the fresh weight of the sample. Statistical analysis was performed using metaboanalyst(60).

### Proteomic analysis

Six weeks old WT plants were grown in short-day conditions. The plants were sprayed with solutions containing no sugars or 0.8mM raffinose (Sigma Aldrich, St. Louis, MO, USA), supplemented with 0.05% tween 20 (J.T Baker, Phillipsburg, NJ, USA). Leaf samples were collected after 24h in 2ml tubes (4 biological replicates per line per treatment). The samples were flash-frozen in liquid nitrogen and stored at −80°C until extraction. Proteins in 5% SDS were precipitated using the chloroform/ methanol method (61). The precipitated proteins were solubilized in 100μl of 8M urea, 10mM DTT, 25mM Tris-HCl pH 8.0 and incubated for 30 min. Iodoacetamide (55mM) was added and followed by incubation for 30 min in the dark. The samples were diluted by the addition of 7 volumes of 25mM Tris-HCl pH 8.0 and sequencing-grade modified Trypsin (Promega Corp., Madison, WI) was added (0.4 μg/ sample) followed by incubation overnight at 37°C with gentle agitation. The samples were acidified by addition of 0.2% formic acid and desalted on C18 home-made Stage tips. Peptide concentration was determined by Absorbance at 280 nm and 0.35 µg of peptides were injected into the mass spectrometer.

MS analysis was performed using a Q Exactive-HF mass spectrometer (Thermo Fisher Scientific, Waltham, MA USA) coupled on-line to a nanoflow UHPLC instrument, Ultimate 3000 Dionex (Thermo Fisher Scientific, Waltham, MA USA). Peptides dissolved in 0.1% formic acid were separated 120 min acetonitrile gradient run at a flow rate of 0.15 μl/min on a reverse phase 25-cm-long C18 column (75 μm ID, 2 μm, 100Å, Thermo PepMapRSLC). Survey scans (300–1,650 m/z, target value 3E6 charges, maximum ion injection time 20 ms) were acquired and followed by higher energy collisional dissociation (HCD) based fragmentation (normalized collision energy 27). A resolution of 60,000 was used for survey scans and up to 15 dynamically chosen most abundant precursor ions, with “peptide preferable” profile were fragmented (isolation window 1.6 m/z). The MS/MS scans were acquired at a resolution of 15,000 (target value 1E5 charges, maximum ion injection times 25 ms). Dynamic exclusion was 20 sec. Data were acquired using Xcalibur software (Thermo Scientific). To avoid a carryover, the column was washed with 80% acetonitrile, 0.1% formic acid for 25 min between samples.

Mass spectra data were processed using the MaxQuant computational platform, version 2.0.3.0. Peak lists were searched against the Uniprot Arabidopsis proteome sequence database (UP000006548) downloaded December 25, 2022 containing 39,324 entries. The search included cysteine carbamidomethylation as a fixed modification, N-terminal acetylation and oxidation of methionine as variable modifications and allowed up to two miscleavages. The match-between-runs option was used. Required FDR was set to 1% at the peptide and protein level. Relative protein quantification in MaxQuant was performed using the label-free quantification (LFQ) algorithm algorithm (62). Statistical analysis was performed using the Perseus statistical (63). Only those proteins for which at least 3 valid LFQ values were obtained in at least one sample group were accepted for statistical analysis. Perseus imputation function was used to replace 0 values. For imputation and volcano plots default Perseus values were used.

## Supporting information

Supporting Figure 1: Raffinose application positively affects plant biomass and in an autophagy-dependent and surfactant-dependent manner.

Supporting Figure 2: Raffinose and galactinol application positively affects plant biomass and in an autophagy-dependent manner.

Supporting Figure 3: Raffinose application positively affects plant growth in several plant species.

Supporting Table S1: Relative polar metabolite content of raffinose-treated plants. Supporting Table S2: Protemoics of raffinose-treated WT and *atg5-1* plants.

## ACKNOWLEDGMENTS

We thank Dr. Hanan Schoffman of the Stein Family Mass Spectrometry Center at the Silberman Institute of Life Sciences, Hebrew University of Jerusalem, for performing the proteomics analysis. We also thank the Avin-Wittenberg group members for their technical assistance and support. The research was funded by the Israel Ministry of Agriculture grant number 12-16-0009.

## DATA AVAILABILITY

All data presented in the manuscript is available in the figures or supplementary material.

## Author Contributions

S.M. designed the research, performed research, analyzed data, and wrote the manuscript. S.D, S.W, D.J.F, S.M., Y.S. and S.M. performed research and analyzed data. T.A-W designed the research and wrote the manuscript.

## Classification

Biological Sciences, Plant Biology

## REFERENCES

1. E. E. Rezaei et al., Climate change impacts on crop yields. Nature Reviews Earth & Environment 4, 831–846 (2023).

2. R. Anderson, P. E. Bayer, D. Edwards, Climate change and the need for agricultural adaptation. Curr Opin Plant Biol 56, 197–202 (2020).

3. T. Avin-Wittenberg, Autophagy and its role in plant abiotic stress management. *Plant*, Cell & Environment 42, 1045–1053 (2019).

4. W. Agbemafle, M. M. Wong, D. C. Bassham, A. Aroca, Transcriptional and post-translational regulation of plant autophagy. Journal of Experimental Botany 74, 6006–6022 (2023).

5. R. S. Marshall, R. D. Vierstra, Autophagy: The Master of Bulk and Selective Recycling. Annual Review of Plant Biology 69, 173–208 (2018).

6. J. H. Doelling, J. M. Walker, E. M. Friedman, A. R. Thompson, R. D. Vierstra, The APG8/12-activating enzyme APG7 is required for proper nutrient recycling and senescence in Arabidopsis thaliana. J Biol Chem 277, 33105–33114 (2002).

7. J. A. S. Barros et al., Autophagy Deficiency Compromises Alternative Pathways of Respiration following Energy Deprivation in Arabidopsis thaliana. Plant Physiology 175, 62–76 (2017).

8. F. Li et al., Autophagic recycling plays a central role in maize nitrogen remobilization. Plant Cell 27, 1389–1408 (2015).

9. S. Ustun, A. Hafren, D. Hofius, Autophagy as a mediator of life and death in plants. Curr Opin Plant Biol 40, 122–130 (2017).

10. E. A. Minina et al., Transcriptional stimulation of rate-limiting components of the autophagic pathway improves plant fitness. J Exp Bot 69, 1415–1432 (2018).

11. T. Xia et al., Heterologous expression of ATG8c from soybean confers tolerance to nitrogen deficiency and increases yield in Arabidopsis. PLoS One 7, e37217 (2012).

12. X. Zhen, F. Xu, W. Zhang, N. Li, X. Li, Overexpression of rice gene OsATG8b confers tolerance to nitrogen starvation and increases yield and nitrogen use efficiency (NUE) in Arabidopsis. Plos One 14 (2019).

13. W.-w. Li et al., Overexpression of the autophagy-related gene SiATG8a from foxtail millet (Setaria italica L.) confers tolerance to both nitrogen starvation and drought stress in Arabidopsis. Biochemical and Biophysical Research Communications 468, 800–806 (2015).

14. E. Baena-Gonzalez, F. Rolland, J. M. Thevelein, J. Sheen, A central integrator of transcription networks in plant stress and energy signalling. Nature 448, 938–942 (2007).

15. J. Soto-Burgos, D. C. Bassham, SnRK1 activates autophagy via the TOR signaling pathway in Arabidopsis thaliana. PLoS One 12, e0182591 (2017).

16. Y. T. Pu, X. J. Luo, D. C. Bassham, TOR-Dependent and -Independent Pathways Regulate Autophagy in Arabidopsis thaliana. Frontiers in Plant Science 8 (2017).

17. L. Chen et al., The AMP-Activated Protein Kinase KIN10 Is Involved in the Regulation of Autophagy in Arabidopsis. Front Plant Sci 8, 1201 (2017).

18. C. M. Figueroa, J. E. Lunn, A Tale of Two Sugars: Trehalose 6-Phosphate and Sucrose. Plant Physiology 172, 7–27 (2016).

19. X. Chen et al., Trehalose, sucrose and raffinose are novel activators of autophagy in human keratinocytes through an mTOR-independent pathway. Scientific Reports 6 (2016).

20. H. C. Janse van Rensburg, W. Van den Ende, S. Signorelli, Autophagy in Plants: Both a Puppet and a Puppet Master of Sugars. Frontiers in Plant Science 10 (2019).

21. B. Williams et al., Trehalose Accumulation Triggers Autophagy during Plant Desiccation. PLoS Genet 11, e1005705 (2015).

22. G. Sun et al., Genome of Paspalum vaginatum and the role of trehalose mediated autophagy in increasing maize biomass. Nat Commun 13, 7731 (2022).

23. S. Alseekh et al., Autophagy modulates the metabolism and growth of tomato fruit during development. Horticulture Research 9 (2022).

24. T. Avin-Wittenberg et al., Global analysis of the role of autophagy in cellular metabolism and energy homeostasis in Arabidopsis seedlings under carbon starvation. Plant Cell 27, 306–322 (2015).

25. S. Sengupta, S. Mukherjee, P. Basak, A. L. Majumder, Significance of galactinol and raffinose family oligosaccharide synthesis in plants. Frontiers in Plant Science 6 (2015).

26. S. Baud, J. P. Boutin, M. Miquel, L. Lepiniec, C. Rochat, An integrated overview of seed development in Arabidopsis thaliana ecotype WS. Plant Physiology and Biochemistry 40, 151–160 (2002).

27. W. V. d. Ende, Multifunctional fructans and raffinose family oligosaccharides. Frontiers in Plant Science 4 (2013).

28. A. Egert, B. Eicher, F. Keller, S. Peters, Evidence for water deficit-induced mass increases of raffinose family oligosaccharides (RFOs) in the leaves of three Craterostigma resurrection plant species. Frontiers in Physiology 6 (2015).

29. A. Egert, F. Keller, S. Peters, Abiotic stress-induced accumulation of raffinose in Arabidopsis leaves is mediated by a single raffinose synthase (RS5, At5g40390). BMC Plant Biology 13 (2013).

30. M. Watanabe et al., Comprehensive dissection of spatiotemporal metabolic shifts in primary, secondary, and lipid metabolism during developmental senescence in Arabidopsis. Plant Physiol 162, 1290–1310 (2013).

31. T. Li et al., Raffinose synthase enhances drought tolerance through raffinose synthesis or galactinol hydrolysis in maize and Arabidopsis plants. Journal of Biological Chemistry 295, 8064–8077 (2020).

32. S. Yan, Q. Liu, W. Li, J. Yan, A. R. Fernie, Raffinose Family Oligosaccharides: Crucial Regulators of Plant Development and Stress Responses. Critical Reviews in Plant Sciences 41, 286–303 (2022).

33. S. Lin, L. Li, M. Li, H. Gu, X. Chen, Raffinose increases autophagy and reduces cell death in UVB-irradiated keratinocytes. Journal of Photochemistry and Photobiology B: Biology 201 (2019).

34. C. Masclaux-Daubresse et al., Stitching together the Multiple Dimensions of Autophagy Using Metabolomics and Transcriptomics Reveals Impacts on Metabolism, Development, and Plant Responses to the Environment in Arabidopsis. Plant Cell 26, 1857–1877 (2014).

35. D. J. Klionsky et al., Guidelines for the use and interpretation of assays for monitoring autophagy (4th edition). Autophagy 10.1080/15548627.2020.1797280, 1–382 (2021).

36. A. Guiboileau et al., Autophagy machinery controls nitrogen remobilization at the whole-plant level under both limiting and ample nitrate conditions in Arabidopsis. New Phytol 194, 732–740 (2012).

37. Y. Pu, J. Soto-Burgos, D. C. Bassham, Regulation of autophagy through SnRK1 and TOR signaling pathways. Plant Signaling & Behavior 12 (2017).

38. G.-H. Chen, M.-J. Liu, Y. Xiong, J. Sheen, S.-H. Wu, TOR and RPS6 transmit light signals to enhance protein translation in deetiolating Arabidopsis seedlings. Proceedings of the National Academy of Sciences 115, 12823–12828 (2018).

39. A. Aroca, I. Yruela, C. Gotor, D. C. Bassham, Persulfidation of ATG18a regulates autophagy under ER stress in Arabidopsis. Proceedings of the National Academy of Sciences 118 (2021).

40. J. A. S. Barros et al., Autophagy during maize endosperm development dampens oxidative stress and promotes mitochondrial clearance. Plant Physiol 193, 1395–1415 (2023).

41. X. Gu, M. Bar-Peled, The Biosynthesis of UDP-Galacturonic Acid in Plants. Functional Cloning and Characterization of Arabidopsis UDP-d-Glucuronic Acid 4-Epimerase. Plant Physiology 136, 4256–4264 (2004).

42. X. Sun et al., MdATG18a overexpression improves tolerance to nitrogen deficiency and regulates anthocyanin accumulation through increased autophagy in transgenic apple. Plant Cell Environ 41, 469–480 (2018).

43. X. Zhen, N. Zheng, J. Yu, C. Bi, F. Xu, Autophagy mediates grain yield and nitrogen stress resistance by modulating nitrogen remobilization in rice. Plos One 16 (2021).

44. T. Avin-Wittenberg et al., Autophagy-related approaches for improving nutrient use efficiency and crop yield protection. Journal of Experimental Botany 69, 1335–1353 (2018).

45. T. Obata et al., Metabolite Profiles of Maize Leaves in Drought, Heat, and Combined Stress Field Trials Reveal the Relationship between Metabolism and Grain Yield. Plant Physiol 169, 2665–2683 (2015).

46. H. Zhang, Y. Zhao, J.-K. Zhu, Thriving under Stress: How Plants Balance Growth and the Stress Response. Developmental Cell 55, 529–543 (2020).

47. E. Baena-González, J. E. Lunn, SnRK1 and trehalose 6-phosphate – two ancient pathways converge to regulate plant metabolism and growth. Current Opinion in Plant Biology 55, 52–59 (2020).

48. N. Reichelt, A. Korte, M. Krischke, M. J. Mueller, D. Maag, Natural variation of warm temperature-induced raffinose accumulation identifies TREHALOSE-6-PHOSPHATE SYNTHASE 1 as a modulator of thermotolerance. Plant, Cell & Environment 46, 3392–3404 (2023).

49. Y. Shin, A. Chane, M. Jung, Y. Lee, Recent Advances in Understanding the Roles of Pectin as an Active Participant in Plant Signaling Networks. Plants 10 (2021).

50. Y. Wang, M. T. Nishimura, T. Zhao, D. Tang, ATG2, an autophagy-related protein, negatively affects powdery mildew resistance and mildew-induced cell death in Arabidopsis. The Plant Journal 68, 74–87 (2011).

51. K. Yoshimoto et al., Autophagy negatively regulates cell death by controlling NPR1-dependent salicylic acid signaling during senescence and the innate immune response in Arabidopsis. Plant Cell 21, 2914–2927 (2009).

52. D. Hofius et al., Autophagic components contribute to hypersensitive cell death in Arabidopsis. Cell 137, 773–783 (2009).

53. A. Mair et al., SnRK1-triggered switch of bZIP63 dimerization mediates the low-energy response in plants. Elife 4 (2015).

54. D. Deprost et al., The Arabidopsis TOR kinase links plant growth, yield, stress resistance and mRNA translation. EMBO reports 8, 864–870 (2007).

55. J. Di Berardino et al., Autophagy controls resource allocation and protein storage accumulation in Arabidopsis seeds. Journal of Experimental Botany 69, 1403–1414 (2018).

56. H. Towbin, T. Staehelin, J. Gordon, Electrophoretic transfer of proteins from polyacrylamide gels to nitrocellulose sheets: procedure and some applications. Proc Natl Acad Sci U S A 76, 4350–4354 (1979).

57. C. A. Schneider, W. S. Rasband, K. W. Eliceiri, NIH Image to ImageJ: 25 years of image analysis. Nat Methods 9, 671–675 (2012).

58. J. Lisec, N. Schauer, J. Kopka, L. Willmitzer, A. R. Fernie, Gas chromatography mass spectrometry-based metabolite profiling in plants. Nat Protoc 1, 387–396 (2006).

59. B. K. Dhatt et al., Metabolic Dynamics of Developing Rice Seeds Under High Night-Time Temperature Stress. Front Plant Sci 10, 1443 (2019).

60. J. Chong, D. S. Wishart, J. Xia, Using MetaboAnalyst 4.0 for Comprehensive and Integrative Metabolomics Data Analysis. Current Protocols in Bioinformatics 68 (2019).

61. D. Wessel, U. I. Flügge, A method for the quantitative recovery of protein in dilute solution in the presence of detergents and lipids. Analytical Biochemistry 138, 141–143 (1984).

62. J. Cox et al., Accurate Proteome-wide Label-free Quantification by Delayed Normalization and Maximal Peptide Ratio Extraction, Termed MaxLFQ. Molecular & Cellular Proteomics 13, 2513–2526 (2014).

63. S. Tyanova et al., The Perseus computational platform for comprehensive analysis of (prote)omics data. Nature Methods 13, 731–740 (2016).

