## Supplementary material for "Raffinose induces autophagy to promote plant growth": Supp. Figures

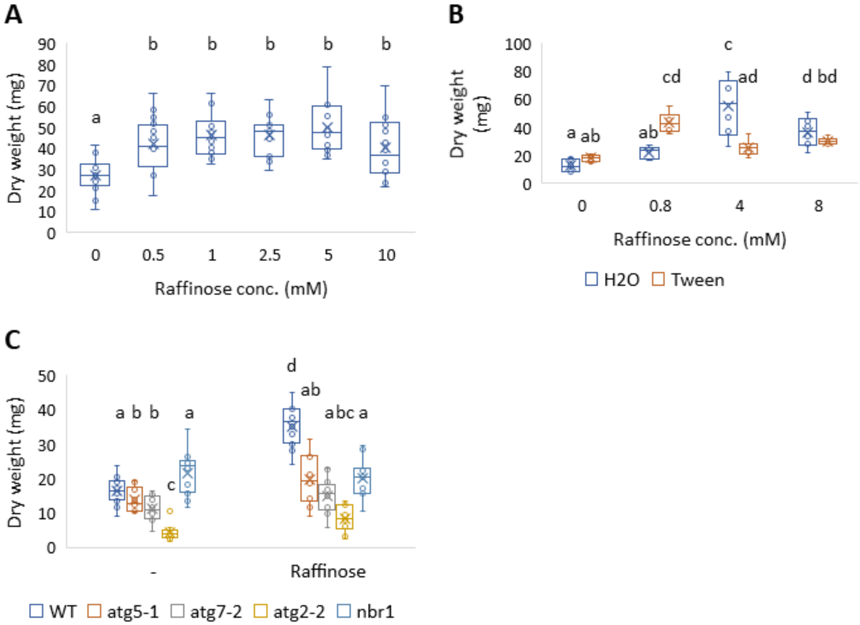


**Supplementary Figure 1: Raffinose application positively affects plant biomass and in an autophagy-dependent and surfactant-dependent manner.** (A) WT Arabidopsis plants were germinated on 1% sucrose Nitsch plates for two weeks and transferred to pots in short-day conditions. After one more week, their foliage was sprayed every week for four weeks with raffinose solution in various concentrations (0,0.5, 1, 2.5, 5,10mM) with 0.05% tween20 as a surfactant. One week after the last spray, entire rosettes were collected, dried and weighed. Data are represented as box&whiskers plot. Letters denote statistical differences between treatments in the WT following ANOVA and Tukey-Kramer test ($p<0.05$, n=13-16). (B) WT and Arabidopsis plants were germinated on 1% sucrose Nitsch plates for two weeks and transferred to pots in short-day conditions. After three more weeks, their foliage was sprayed every two weeks with raffinose solution in various concentrations (0,0.8,4,8mM) with or without 0.05% tween20 as a surfactant. One week after the second spray, entire rosettes were collected, dried and weighed. Data are represented as box&whiskers plot. Letters denote statistical differences between treatments in the WT following ANOVA and Tukey-Kramer test ($p<0.05$, n=12). (C) WT, *atg5-1, atg7-2, atg2-2 and nbr1* mutant Arabidopsis plants were germinated on 1% sucrose Nitsch plates for two weeks and transferred to pots in short-day conditions. After one more week, their foliage was sprayed every week for four weeks with raffinose solution in two concentrations (-: 0, raffinose: 1mM) with 0.05% tween20. One week after the last spray, entire rosettes were collected, dried and weighed. Letters denote statistical differences between treatments in the WT following ANOVA and Tukey-Kramer test ($p<0.05$, n=16).

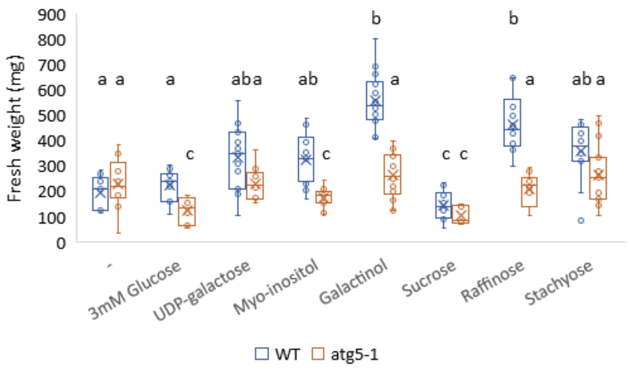


**Supplementary Figure 2: Raffinose and galactinol application positively affects plant biomass and in an autophagy-dependent manner.** WT and *atg5-1* Arabidopsis plants were germinated on 1% sucrose Nitsch plates for two weeks and transferred to pots in short-day conditions. After one more week, their foliage was sprayed every week for four weeks with solutions containing so sugars (-), 1mM raffinose precursors or 3mM glucose with 0.05% tween20 as a surfactant. One week after the last spray, entire rosettes were collected and  rosette fresh weight measured. Data are represented as box&whiskers plot. Letters denote statistical differences between treatments in the WT following ANOVA and Tukey-Kramer test (p<0.05, n=5-10).

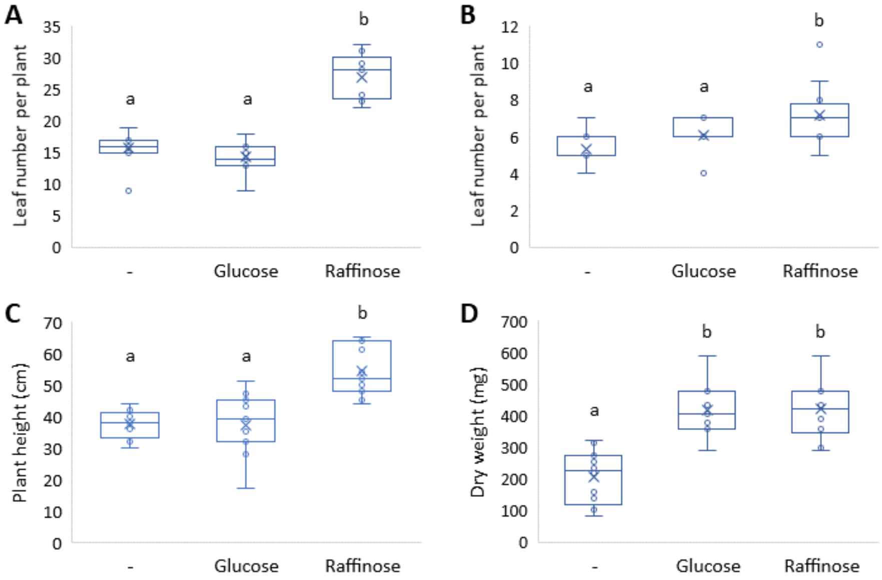


**Supplementary Figure 3: Raffinose application positively affects plant growth in several plant species.** (A) Tobacco (*Nicotiana benthamiana*) plants were grown in long-day conditions for 3 weeks. The plants were sprayed with solutions containing no sugars (-), 0.8mM raffinose or 3mM glucose, supplemented with 0.05% tween20 once a week for three weeks. Leaf number per plant was scored one week after the third spraying. Data are presented as box&whiskers plot. Letters denote statistically significant treatments following ANOVA and post-hoc Tuckey-Kramer test ($p<0.05$, n=9). B-D. Wheat *(Triticum aestivum*) plants were grown in long-day conditions for 2 weeks. The plants were sprayed with solutions containing no sugars (-), 0.8mM raffinose, or 3mM glucose, supplemented with 0.05% tween20 once a week for three weeks. B. Leaf number per plant one week after the third spraying. C. Plant height one week after the third spraying. D. Foliage dry weight one week after the third spraying. Data are presented as box&whiskers plot. Letters denote statistically significant treatments following ANOVA and post-hoc Tuckey-Kramer test ($p<0.05$, n=11).
